# Identification of a prostaglandin E2 receptor that regulates mosquito oenocytoid immune cell function in limiting bacteria and parasite infection

**DOI:** 10.1101/2020.08.03.235432

**Authors:** Hyeogsun Kwon, David R. Hall, Ryan C. Smith

**Affiliations:** Department of Entomology, Iowa State University, Ames, Iowa 50010, USA

## Abstract

Lipid-derived signaling molecules known as eicosanoids have integral roles in mediating immune and inflammatory processes across metazoans. This includes the function of prostaglandins and their cognate G protein-coupled receptors (GPCRs) to employ their immunological actions. In insects, prostaglandins have been implicated in the regulation of both cellular and humoral immune responses, yet studies have been limited by the absence of a described prostaglandin receptor. Here, we characterize a prostaglandin E2 receptor (*Ag*PGE2R) in the mosquito *Anopheles gambiae* and examine its contributions to innate immunity. *Ag*PGE2R expression is most abundant in circulating hemocytes where it is primarily localized to oenocytoid immune cell populations. Through the administration of prostaglandin E2 (PGE2) and *AgPGE2R*-silencing by RNAi, we demonstrate that PGE2 signaling regulates the expression of a subset of prophenoloxidases (PPOs) and antimicrobial peptides (AMPs). PGE2 priming via the *Ag*PGE2R significantly limited bacterial replication and suppressed *Plasmodium* oocyst survival. Additional experiments establish that PGE2 priming increases phenoloxidase (PO) activity through the increased expression of *PPO1* and *PPO3*, which significantly influence *Plasmodium* oocyst survival. We also provide evidence that PGE2 priming is concentration-dependent, where high concentrations of PGE2 promote oenocytoid lysis, negating the protective effects of PGE2 priming on anti-*Plasmodium* immunity. Taken together, our results characterize the *Ag*PGE2R and the role of prostaglandin signaling on immune cell function, providing new insights into the role of PGE2 on anti-bacterial and anti-*Plasmodium* immune responses in the mosquito host.

## Introduction

Eicosanoids are lipid-derived signaling molecules that include prostaglandins (PGs), leukotrienes (LTs), lipoxins (LXAs), and epoxyeicosatrienoic acid (EETs), that serve important roles in immune regulation (1–3). Evidence suggests that these responses are evolutionally conserved across metazoa, where eicosanoids significantly influence insect cellular immunity (4–10). In the mosquito, *Anopheles gambiae*, eicosanoids such as prostaglandin E2 (PGE2) and lipoxins have integral roles in mediating immune priming and mosquito susceptibility to malaria parasite infection (11, 12). However, our understanding of eicosanoid-mediated immune regulation in insects has remained incomplete due to the lack of characterized eicosanoid biosynthesis pathways and cognate receptors to initiate signaling. Therefore, the identification and characterization of eicosanoid receptors are needed to properly examine the roles of eicosanoids in insect immunity.

The recent characterization of a PGE2 receptor in the lepidopteran systems *Manduca sexta* (8) and *Spodoptera exigua* (10) have implicated the PGE2 receptor in immune activation pathogen-associated molecular patterns (PAMPs)(8) and cellular immune function (13). Yet, despite the implication of PGE2 on mosquito immunity (4, 11), the cognate prostaglandin receptor has not yet been described in mosquitoes. Therefore, the aim of this study was to identify and characterize the PGE2 receptor (*Ag*PGE2R) in the mosquito *An. gambiae*, and to describe its function in the mosquito innate immune response.

Based on *in silico* analyses using the *Manduca* PGE2 receptor (8) and human PG receptors, we identified a putative PGE2 receptor (*Ag*PGE2R) in *An. gambiae* orthologous to human PGE2-EP2 and EP4 receptors. Herein, we provide compelling evidence that *Ag*PGE2R is predominantly expressed in the oenocytoid immune cell populations and promotes the production of antimicrobial peptides (AMPs) and a subset of prophenoloxidases (PPOs) that limit bacterial and *Plasmodium* survival in the mosquito host. Therefore, our study provides important new insights into mosquito PGE2 signaling, oenocytoid cell function, and the immune mechanisms that limit pathogens in the mosquito host. Moreover, our study represents a significant advance in our understanding of prostaglandin signaling in insect innate immunity, as well as an invaluable resource for comparative immunology to identify eicosanoid receptors in other dipteran species.

## Results

### Characterization of the *Ag*PGE2 receptor

Based on the prostaglandin E2 receptor from the tobacco hornworm, *Manduca sexta*(8), a putative PGE2 receptor (*Ag*PGE2R; AGAP001561) was identified in *An. gambiae*. Based on this annotation, a full-length transcript consisting of 1463 bp and encoding a 404 amino acid protein (∼46 kDa) was isolated from perfused naïve *An. gambiae* hemolymph (Figure S1). *In silico* analysis supports that the *Ag*PGE2R contains 7 transmembrane domains and belongs to the rhodopsin-like family of G protein-coupled receptors (GPCRs; Figures S1 and S2). Moreover, four N-glycosylated and six phosphorylation sites are predicted at the respective N- and C-terminus (Figures S1 and S2A), suggesting that the *Ag*PGE2R undergoes post-translational modifications as in mammalian systems (14, 15). *Ag*PGE2R also contains core residues characteristic of other Family A GPCRs and human prostanoid receptors (Figure S2A). Phylogenetic analysis reveals a unique clade of insect receptors where *Ag*PGE2R is most similar to other putative dipteran PGE2 receptors (Figure S2B). In addition, insect PGE2Rs are more closely related to human prostanoid receptors than human leukotriene receptors (CYSLTR1/2), which also serve as important eicosanoid receptors belonging to Family A GPCRs (Figure S2B).

To more closely characterize the role of the putative mosquito prostaglandin receptor, we examined *AgPGE2R* expression in multiple tissues from different physiological conditions (naïve, 24 h blood-fed, and 24 h *P. berghei<u>-</u>*infected; Figure 1A). Under naïve conditions, receptor expression is highly enriched in hemocytes (Figure 1A). In addition, *AgPGE2R* was also highly expressed in the malpighian tubules following blood-feeding, suggesting potential roles for the receptor in maintaining water homeostasis after blood-feeding (Figure 1A). However, when receptor expression was evaluated in the same tissue under different feeding conditions, only hemocytes displayed significant differences in *AgPGE2R* expression (Figure 1B, Figure S3). Western blot analysis using mosquito hemolymph samples result in a single band of approximately 70 kDa (Figure 1C), much higher than the expected ∼46 kDa MW of the annotated *Ag*PGE2R. Enzymatic deglycosylation through the use of PNGase F decreased the size of the *Ag*PGE2R protein to the expected size of ∼46 kDa (Figure 1C), indicating that the *Ag*PGE2R undergoes substantial post-translational modifications by glycosylation. No specific bands were detected with the incubation of pre-immune serum (Figure 1C). Immunofluorescence assays revealed that *Ag*PGE2R is predominantly expressed in oenocytoid immune cell populations (Figure 1D), although weak signal is also detected in a sub-set of granulocytes (Figure S4A). To further validate *Ag*PGE2R localization to the oenocytoid, phagocytic granulocyte populations were ablated by clodronate liposome (CLD) treatment as previously described (16). qRT-PCR analysis demonstrates that *Ag*PGE2R transcript remains unchanged in perfused hemolymph samples from mosquitoes treated with CLD, while the expression of *eater* (a known phagocytic maker (17, 18)) was significantly reduced (Figure S4B). Taken together, these results support that *Ag*PGE2R is expressed primarily in oenocytoid immune cell populations and confirmed by recent scRNA-seq studies of mosquito immune cell populations (19).

**Figure 1.**
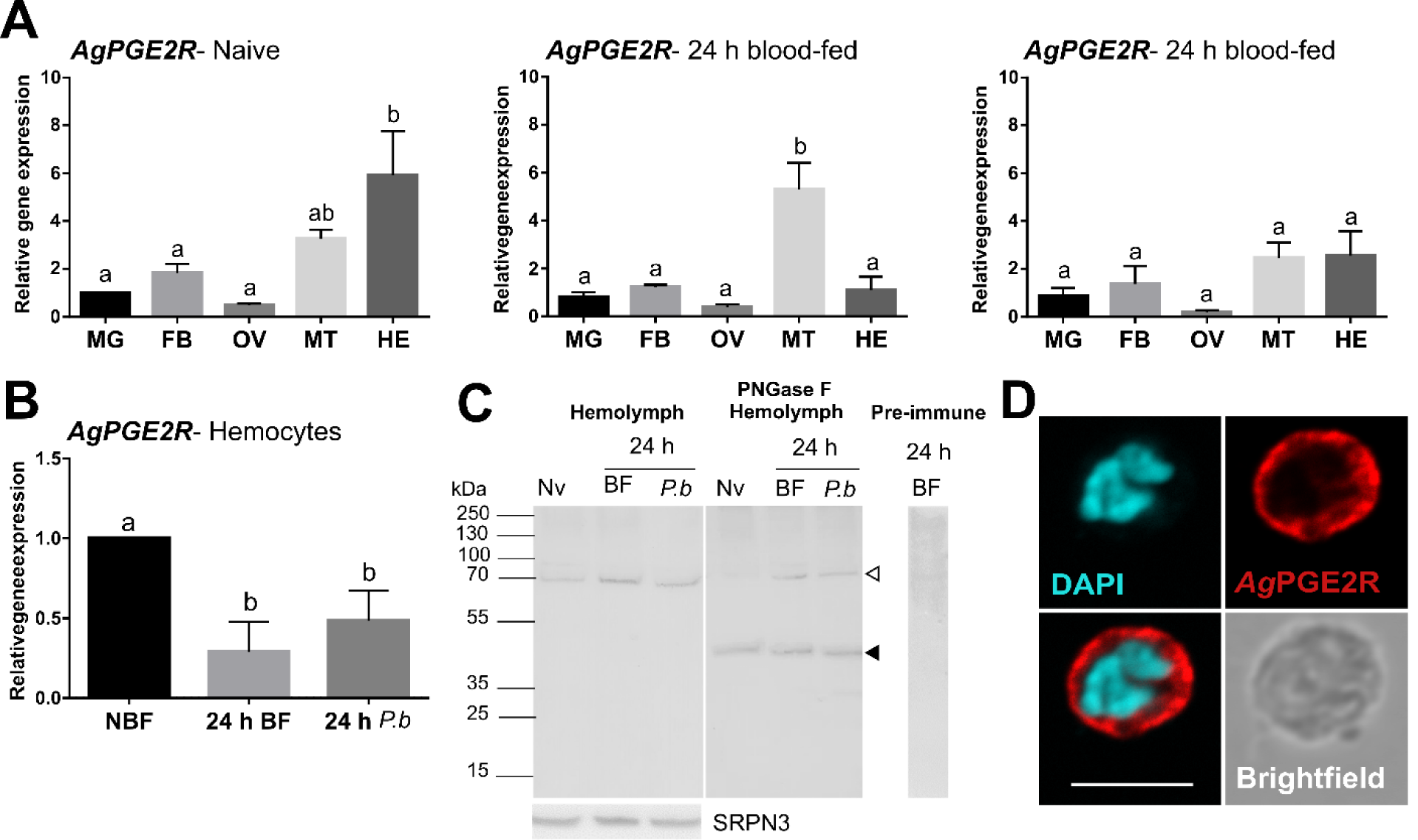
Characterization of a prostaglandin E2 receptor (PGE2R) in *Anopheles gambiae*. (**A**) *Ag*PGE2R gene expression was examined by qRT-PCR in midgut (MG), fat body (FB), ovary (OV), malphigian tubules (MT), and perfused hemocytes (HE) under naïve (NV), blood-fed (24 h BF), or *P. berghei*-infected (24 h *P*.*b*) conditions. (**B**) *Ag*PGE2R expression was more closely examined in hemocytes and compared across physiological conditions. For both **A** and **B**, letters indicate statistically significant differences (*P* < 0.05) when analyzed by a one-way ANOVA followed by a Tukey’s multiple comparison test using GraphPad Prism 6.0. Bars represent the mean ± SE of either three or four independent biological replicates. (**C**) Perfused hemolymph from naïve (NV), blood-fed (BF), or *P. berghei* infected (*P*.*b*) mosquitoes were examined by Western blot analysis. Specific bands were detected corresponding to a glycosylated *Ag*PGE2R product (bands at ∼70 kDa, open arrowhead), or in which the receptor underwent deglycosylation with PNGase F treatment (∼46 kDa, black arrowhead). The *Ag*PGE2R was detected using a rabbit antibody (1:1000) directed against intracellular loop 3 (ICL3). No bands were detected in a 24 h BF hemolymph sample incubated with pre-immune serum. Serpin 3 (SRPN3) was used as a protein loading control. (**D**) Immunofluorescence assays were performed on perfused hemocytes using a rabbit polyclonal antibody (1:500) against extracellular loop 2 (ECL2), revealing that PGE2R is localized to oenocytoid immune cell populations (scale bar, 5 µm).

### PGE2 biosynthesis and the influence of PGE2 signaling on mosquito immunity

To understand the molecular mechanisms of PGE2 signaling in *An. gambiae*, PGE2 titers were measured from whole mosquitoes or perfused hemolymph under naïve, 24 h blood-fed, or 24 h *P. berghei-*infected conditions. In whole mosquitoes, PGE2 titers were significantly increased in blood-fed and infected mosquitoes when compared to naïve mosquitoes (Figure S5). Yet, similar to previous reports(11), PGE2 levels in the hemolymph reach measurable levels only following *P. berghei* infection (Figure S5). These results suggest that the ELISA assay may detect PGE2 present in mouse blood (20), while PGE2 levels in the hemolymph are increased in response to ookinete invasion (11). Based on our earlier observations of *Ag*PGE2R localization in oenocytoid cell populations (Figure 1D), we explored what role PGE2 signaling may have on prophenoloxidase (PPO) expression. qRT-PCR analysis demonstrates that a subset of PPOs which includes *PPO1, PPO3, PPO7* and *PPO8* were upregulated in response to PGE2 priming (Figure 2A). In contrast, *AgPGE2R*-silencing (Figure S6) downregulated the same subset of *PPOs* (Figure 2B), suggesting that *PPO1, PPO3, PPO7* and *PPO8* are regulated by PGE2 signaling. To determine if the PGE2-mediated PPO regulation also influences phenoloxidase (PO) activity, we performed dopa conversion assays in response to PGE2 signaling. Compared to PBS controls, PGE2 priming significantly increased hemolymph PO activity (Figure 2C). Moreover, PGE2 priming in *Ag*PGE2R-silenced mosquitoes resulted in substantially less PO activity than GFP-silenced controls (Figure 2D), providing additional support that PGE2 signaling promotes PO activity via the *Ag*PGE2R.

**Figure 2.**
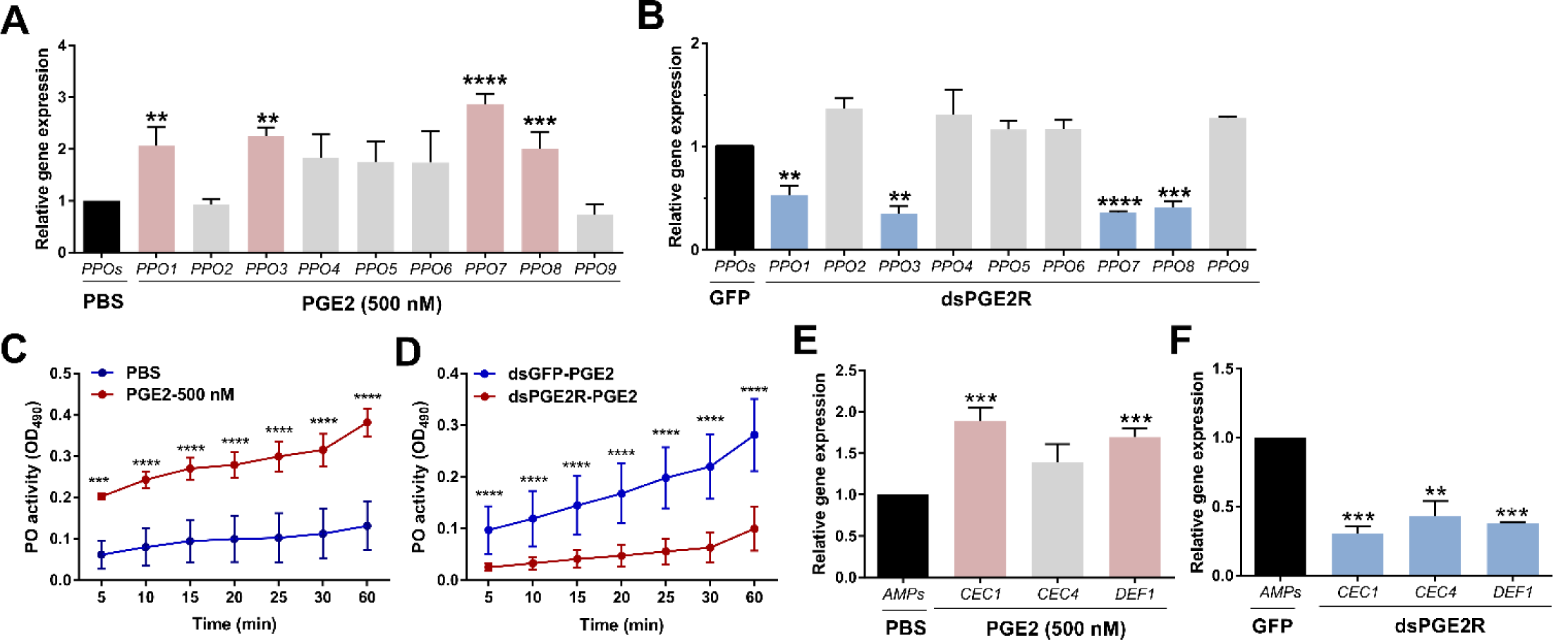
PGE2 signaling influences phenoloxidase (PO) activity and antimicrobial peptide (AMP) gene expression. Following treatment with PGE2 (500nM) (**A**) or when *AgPGE2R* was silenced (**B**), the expression of all 9 mosquito prophenoloxidases (PPOs) were examined by qRT-PCR and compared to respective PBS or GFP controls. For both **A** and **B**, mosquitoes were pooled (n=15) for analysis. Data were analyzed using an unpaired t test to determine differences in relative gene expression for each respective PPO gene between treatments. Bars represent mean ± SE of three independent biological replicates. (**C**) PO activity was measured from perfused hemolymph in mosquitoes primed with PGE2 (500 nM) and compared to PBS controls 24 h post-treatment (n=15 per treatment). (**D**) Additional experiments were performed measuring PO activity in which PGE2 (500nM) was injected into *AgPGE2R*- or *GFP*-silenced (control) mosquitoes (n=15 per treatment). For both **C** and **D**, measurements (OD490) were taken for DOPA conversion assays at 5-min intervals from 0 to 30 min, as well as a final readout at 60 min. Data were analyzed using a two-way repeated-measures ANOVA followed by Sidak’s multiple comparisons using GraphPad Prism 6.0. Bars represent mean ± SE of 3 independent experiments. In addition, PGE2 priming induced expression of antimicrobial peptide (AMP) genes (**E**), while *AgPGE2R-silencing* reduced AMP expression (**F**). For **E** and **F**, data were analyzed using an unpaired *t*-test to determine differences in relative gene expression of each respective AMP gene between treatments. Bars represent mean ± SE of three independent replications. For all data, asterisks denote significance (\*\**P* < 0.01, \*\*\**P* < 0.001, \*\*\*\**P* < 0.0001); ns, not significant.

In addition, previous studies have implicated PGE2 signaling on the synthesis of antimicrobial peptides (AMPs) in *Anopheles albimanus* (4). Therefore, we examined the expression of major AMPs in mosquitoes primed with PGE2 or following *Ag*PGE2R-silencing. Similar to the regulation of PPOs, PGE2 priming significantly increased the expression of *cecropin 1* (*CEC1*) and *defensin 1* (*DEF1*) (Figure 2E), while the silencing of *Ag*PGE2R reduced *CEC1, CEC4* and *DEF1* (Figure 2F). These data support that prostaglandin signaling via *Ag*PGE2R, directly influences PPO expression and subsequent PO activity, as well as AMP expression.

### PGE2 signaling promotes antimicrobial activity

Based on the regulation of AMP expression in response to PGE2 treatment and requiring *AgPGE2R* function (Figure 2), we examined the antimicrobial effects of PGE2 signaling by challenging mosquitoes with *E. coli* following PGE2 priming. At 6 h post-challenge, PGE2-primed mosquitioes displayed significantly less bacteria when compared to PBS controls (Figure S7A). We identified a similar trend at 24 h post-challenge, yet these results were not significant (Figure S7A). In addition, we demonstrate that the effects of PGE2 priming are abrogated following *AgPGE2R*-silencing (Figure S7B), providing further support that prostaglandin signaling limits bacterial growth

### PGE2 signaling mediates anti-*Plasmodium* immunity

In agreement with the previously reported effects of PGE2 priming on anti-*Plasmodium* immunity (11), we confirm that mosquitoes primed with PGE2 have significantly reduced oocyst numbers eight days post-infection (Figure 3A). To confirm the involvement of *Ag*PGE2R in mediating these responses, we performed RNAi experiments to silence the receptor and evaluate its contributions to *Plasmodium* survival. As previously described, RNAi significantly reduced *AgPGE2R* at four days post-injection, as well as 24 h post-*P. berghei* infection (Figure S6). *AgPGE2R*-silencing had no effect on early oocyst numbers two days post-infection, yet resulted in an increase in oocyst survival at eight days post-infection (Figure 3B) similar to other known mediators of mosquito late-phase immunity (16, 21–24). To further validate the role of *Ag*PGE2R in mediating the anti-*Plasmodium* effects of PGE2 signaling, we demonstrate that PGE2 priming requires *Ag*PGE2R function (Figure 3C).

**Figure 3.**
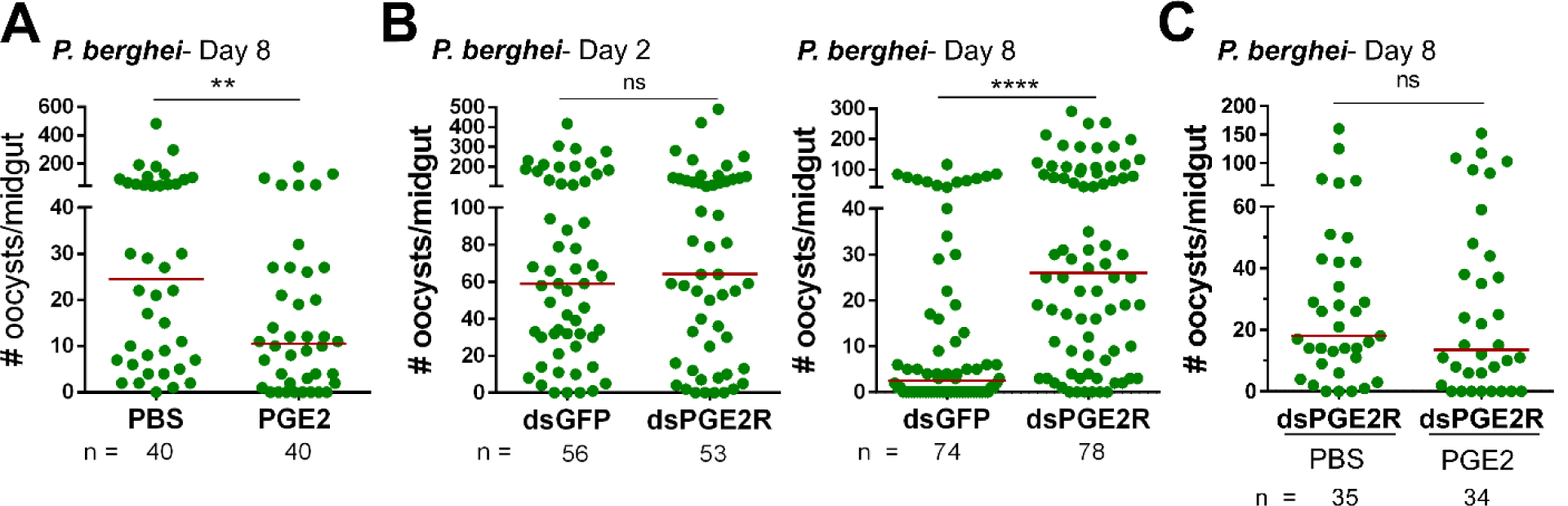
PGE2 signaling mediates *An. gambiae* anti-*Plasmodium* immunity. Following priming with PGE2 (500 nM), *P. berghei* infection was measured by evaluating oocyst numbers 8 days post-infection (**A**) and compared to PBS controls. When *AgPGE2R*-silenced mosquitoes were challenged with *P. berghei*, early oocyst numbers (Day 2) were comparable to GFP controls, yet displayed significant differences at Day 8 (**B**), reminiscent of previously described late-phase immunity phenotypes. When PGE2 was administered to *AgPGE2R*-silenced mosquitoes, there were no effects of PGE2 priming on oocyst survival (**C**). All infection data were analyzed by a Mann–Whitney test using GraphPad Prism 6.0. Median oocyst numbers are indicated by the horizontal red line. Bars represent mean ± SE of three or more independent biological replicates. Asterisks denote significance (\*\**P* < 0.01, \*\*\*\**P* < 0.0001); ns, not significant.

Since PGE2 signaling regulates the expression of multiple PPO and AMP genes (Figure 2), we hypothesized that these immune molecules may directly mediate the effects of PGE2 priming on oocyst survival. Previous studies have shown that PPO3 (16) and CEC3 (25) antagonize *Plasmodium* survival, while DEF1 does not directly impact parasite infection (26). However, *PPO-1, 7, 8* and *CEC1* have not previously been evaluated by RNAi for their role in anti-*Plasmodium* immunity. While each transcript was significantly reduced (Figures 4A and S8A), only PPO1 significantly influenced oocyst survival at seven days post-infection (Figures 4A). With previous evidence supporting that PPOs contribute to the late-phase immune responses that mediate oocyst survival(16), we wanted to examine the potential that PPO1 and PPO3 were responsible for the anti-*Plasmodium* effects of PGE2 signaling. When *PPO1* and *PPO3* were co-silenced (Figure 4B), PGE2 priming had no effect on *Plasmodium* oocyst numbers at seven days post-infection, in contrast to GFP-silenced controls (Figure 4B). This provides strong support that PPO1 and PPO3 mediate the oocyst killing responses produced by PGE2 signaling activation.

**Figure 4.**
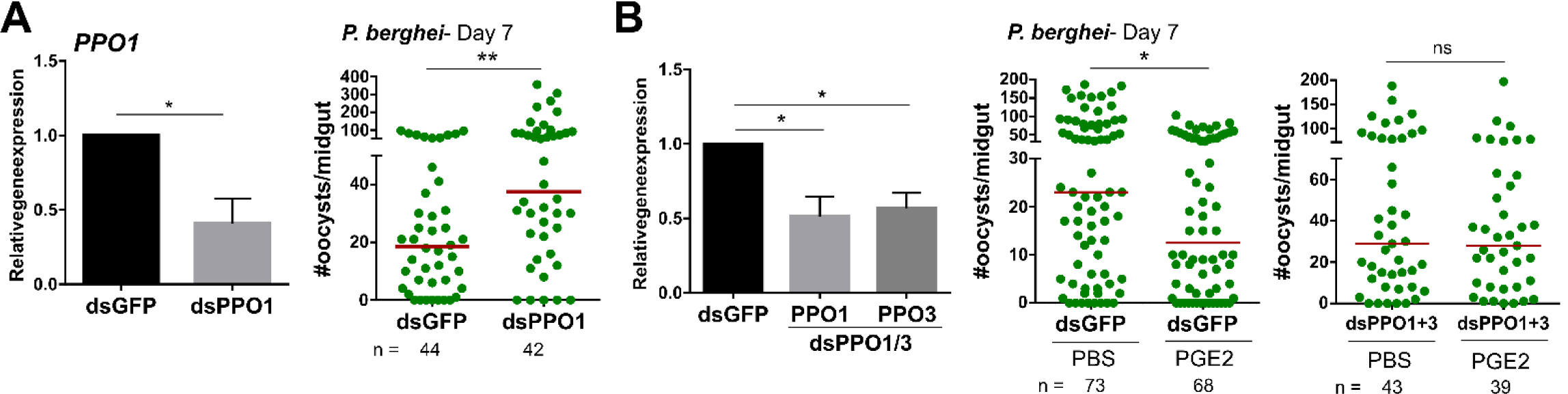
Anti-*Plasmodium* effects of PGE2 priming are mediated by mosquito prophenoloxidases. Significant reduction of PPO1 gene expression enhanced oocyst survival at 7 days post-infection (**A**) *PPO1* gene expression was efficiently reduced following RNAi, resulting in a significant increase in P. berghei oocyst survival when evaluated 8 days post-infection. To verify the role of mosquito prophenoloxidases in mediating PGE2 immune activation, *PPO1* and *PPO3* were co-silenced and then primed with either PBS (control) or PGE2 (**B**). Following challenge with *P. berghei*, oocyst survival was examined 7 days post-infection. Data were analyzed by an unpaired t test to determine RNAi efficiency and a Mann–Whitney test to assess oocyst survival using GraphPad Prism 6.0. Bars represent mean ± SE of three independent biological replicates. Median oocyst numbers are indicated by the horizontal red line. Asterisks denote significance (\**P* < 0.05, \*\**P* < 0.01); ns, not significant.

### PGE2 triggers oenocytoid cell lysis

Given the increase in PO activity following PGE2 treatment (Figure 2), and studies implicating PGE2 in the release of PPOs as a result of oenocytoid cell lysis in lepidopteran insects (6, 27), we investigated whether PGE2 might similarly induce oenocytoid lysis in *An. gambiae*. In contrast to the induction of *PPO* genes upregulated by PGE2 treatment at 500 nM (Figure 2A), higher concentrations of PGE2 (1 µM and 2 µM) decreased the expression of the same subset of PPO genes as compared to PBS controls (Figures S10 and 5A). Moreover, relative *AgPGE2R* transcript was similarly reduced in mosquitoes primed with higher PGE2 concentrations (Figures S10 and 5B), although *AgPGE2R* expression was unchanged by PGE2 priming at 500 nM (Figure 5B). To more closely examine these differences, we focused on only the 500 nM and 2 µM PGE2 concentrations. Using immunofluorescence assays (IFAs), we demonstrate that the administration of 500 nM of PGE2 has no effect on the proportion of *Ag*PGE2R^+^ cells, yet at higher concentrations of PGE2 (2 µM), the proportion of *Ag*PGE2R^+^ cells was significantly reduced (Figure 5C) with a corresponding increase in PO activity (Figure S9). In intact oenocytoid cells, transcripts such as *AgPGE2R* and *PPO1* can readily be distinguished, yet can no longer be detected after cell lysis (Figure 5). Together, these results support that high concentrations of PGE2 initiate oenocytoid lysis, releasing their contents into the mosquito hemolymph that initiate subsequent PO activity. To determine the effects of oenocytoid lysis on malaria parasite survival, mosquitoes pretreated with 2 µM of PGE2 were challenged with *P. berghei* infection. Interestingly, when mosquitoes were primed with 2 µM of PGE2 to reduce oenocytoid populations before *P. berghei* challenge, oocyst survival was significantly increased (Figure 5D), suggesting that oenocytoid populations are integral to anti-*Plasmodium* immunity and the effects of PGE2 priming.

**Figure 5.**
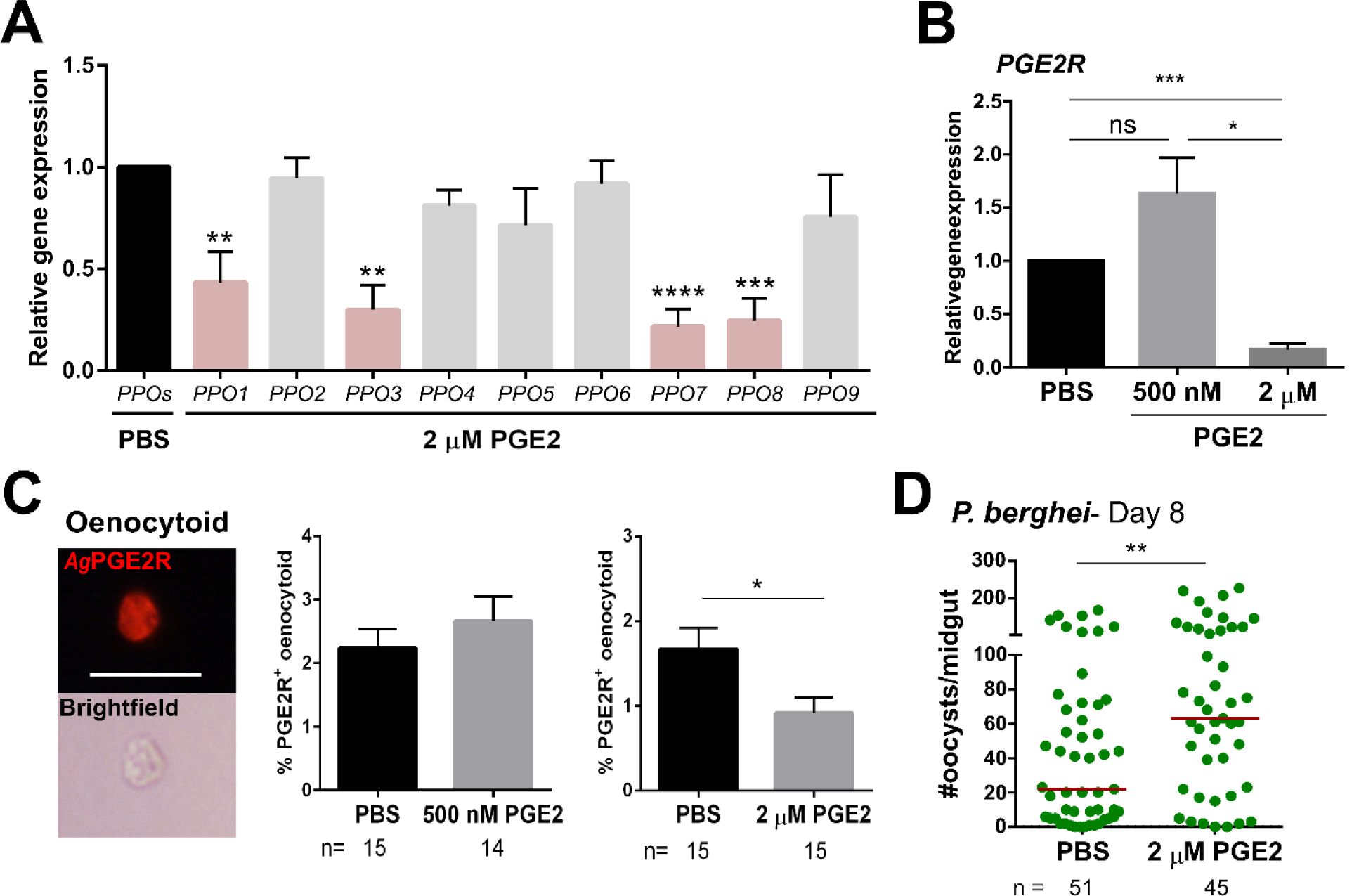
High concentrations of PGE2 trigger oenocytoid lysis. (**A**) The influence of high PGE2 concentrations (2 µM) was examined on prophenoloxidase (PPO) gene expression, where the subset of PPO genes regulated by PGE2 treatment (*PPO1, 3, 7* and *8*) are significantly reduced. (**B**) Similarly, *AgPGE2R* expression was also reduced when mosquitoes were primed with 2 µM PGE2, yet was unaffected by 500 nM PGE2. To determine if oenocytoid immune cell populations were influenced by PGE2 levels, immunofluorescence assays (IFAs) were performed to label mosquito oenocytoids (**C**). There was no difference in the proportion of *Ag*PGE2R+ oenocytoid cells between PBS and 500 nM PGE2 treatments, yet 2 µM PGE2 treatments reduced oenocytoid populations supporting that high levels of PGE2 promote oenocytoid lysis. The scale bar represents 10 µm. (**D**) When oenocytoid lysis was triggered by 2 µM PGE2 treatment prior to *P. berghei* infection, *Plasmodium* survival was significantly increased when oocyst numbers were examined 8 days post-infection. Data presented in **A**-**C** were analyzed using an unpaired t-test to determine differences in relative gene expression or oenocytoid abundance using GraphPad Prism 6.0. Bars represent mean ± SE of three independent biological replicates for **A** and **B**, and two independent biological replicates for **C**. Oocyst data from three independent experiments were analyzed by Mann–Whitney. Median oocyst numbers are indicated by the horizontal red line. Asterisks denote significance (\**P* < 0.05, \*\**P* < 0.01, \*\*\**P* < 0.001, \*\*\*\**P* < 0.0001); ns, not significant.

## Discussion

The interactions between the innate immune system and malaria parasites are key determinants in shaping mosquito vector competence (28, 29). Therefore, much effort has been devoted to define the immune molecules and immunological mechanisms that determine malaria parasite killing in the mosquito host. Over the last 30 years, prostaglandins have been implicated in mediating insect cellular immune responses such as hemocyte spreading and chemotaxis, nodule formation, melanization, and encapsulation (11, 13, 30, 31). However, our current understanding of the immunological mechanisms of prostaglandin signaling have been limited by the lack of characterized prostaglandin receptors.

In this study, we cloned and sequenced a putative PGE2 receptor (*Ag*PGE2R) in *An. gambiae*, exhibiting similarities to recently characterized insect PGE2 receptors in *M. sexta* (8) and *S. exigua* (13). With prostaglandin signaling implicated in diverse physiological roles in insects such as cellular immunity, renal physiology and reproduction (1, 8, 11, 32–34), we examined *AgPGE2R* expression across tissues and feeding status, finding that *AgPGE2R* was most prevalent in populations of hemocytes and malphigian tubules. While we did not further explore the role of *AgPGE2R* in the malpighian tubule, previous studies in other insect systems suggest potential roles for *Ag*PGE2R in water homeostasis (33, 35, 36). With significant roles described for prostaglandins in the insect cellular immune response (6, 8, 11), our efforts herein are focused on the function of *AgPGE2R* in mosquito immune cell populations where the receptor was most highly expressed.

Using immunofluorescence assays, we demonstrate that *Ag*PGE2R is predominantly expressed in the oenocytoid, a non-phagocytic immune cell sub-type traditionally associated with prophenoloxidase (PPO) production (37). However, *Ag*PGE2R was also immunolocalized in a subset of phagocytic cells at a lower signal intensity. These cell identifications are supported herein by the use of clodronate liposomes (16) to demonstrate that *AgPGE2R* is unaffected by phagocyte depletion experiments, and recent single-cell RNA-seq studies of *An. gambiae* immune cell populations (19). Our findings are also consistent with recent evidence that the *M. sexta* PGE2R is immunolocalized to oenocytoid immune cell populations (8). Based on these expression patterns, we therefore argue that oenocytoids are central to prostaglandin signaling.

Our data demonstrate an integral role for *Ag*PGE2R in oenocytoid function, influencing the transcription of PPO and AMP genes, PO activity, and oenocytoid lysis. Moreover, these data provide definitive evidence for the role of oenocytoids in mosquito innate immunity, with involvement in limiting bacteria and malaria parasite survival (summarized in Figure 6). However, we cannot discount other PGE2 signaling events that may influence the immune system. With weak expression on a subset of granulocytes, *Ag*PGE2R may also directly or indirectly mediate hemocyte recruitment to the midgut basal lamina and granulocyte proliferation as previously described (11). In addition, the expression of *AgPGE2R* in other tissues, such as the salivary gland, may also have important roles in mosquito physiology that require further investigation.

**Figure 6.**
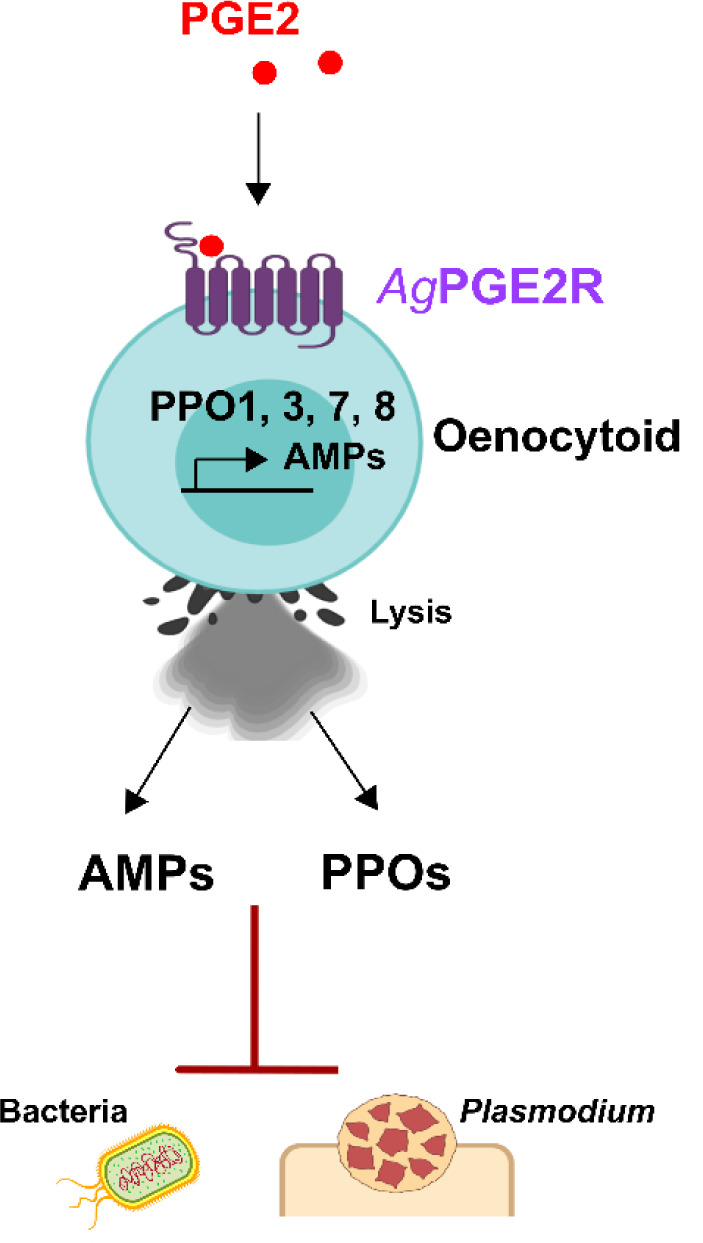
Summary of PGE2 signaling via PGE2R to activate *A. gambiae* immunity in oenocytoid immune cell populations. We demonstrate that PGE2 interacts with its cognate receptor, *Ag*PGE2R, to initiate innate immune expression of prophenoloxidases and AMPs in oenocytoid immune cell populations. The release of these molecules via ookinete lysis mediate the killing of bacteria and malaria parasites.

Focusing on the influence of *Ag*PGE2R in mosquito immune function, we demonstrate that PGE2 priming influences PPO gene expression and subsequent downstream PO activity. Of interest, we found that only a subset of PPO genes (*PPO1, -3, -7* and *-8*) were influenced by prostaglandin signaling. This same subset of PPO genes was upregulated in response to PGE2 priming, while *AgPGE2R*-silencing significantly reduced their expression. As a result, these data support that prostaglandin signaling is central to the regulation of *PPO1, -3, -7* and *-8*. This same subset of PPOs is highly enriched in mosquito oenocytoid popoulations and display regulation by the transcription factor lozenge(19), an important regulator of oenocytoid (or comparable *Drosophila* crystal cell) differentiation(19, 38, 39). Recent data suggest that the remaining PPO genes, *PPO2*, - *4*, -*5*, -*6*, -*9* are highly expressed in granulocyte populations (16, 19), providing additional support for distinct mechanisms of PPO regulation in mosquito immune cell populations.

Previous studies have demonstrated that injury and bacterial infection cause the release of PPO from *Drosophila* crystal cells (40), supporting that the increase in PGE2 levels that accompany ookinete invasion may similarly trigger oenocytoid lysis. We provide evidence that PGE2 promotes oenocytoid lysis in a concentration-dependent manner, where increasing PGE2 concentrations result in increased oenocytoid lysis. This is supported by the role of prostaglandins in mediating the release of PPO through oenocytoid rupture in the beet armyworm *S. exigua* (6, 27, 41), although evidence suggests that lysis is independent of the prostaglandin receptor (6, 41).

Our data support that PPO regulation via prostaglandin signaling is an integral component of mosquito “late-phase” immunity that limits *Plasmodium* oocyst survival (16, 21–23, 28). Similar to the phenotypes previously described for *PPO2*-, *PPO3*- and *PPO9*-silencing (16), we demonstrate that *PPO1*-silencing increase the post-invasion survival of immature oocyts. In addition, co-silencing of *PPO1* and *PPO3* eliminated the protective effects of PGE2 priming, arguing that PPO expression is central to anti-*Plasmodium* effects of prostaglandin signaling. This is further supported by our experiments using high PGE2 concentrations to deplete oenocytoid popoulations prior to *P. berghei* challenge which importantly implicate oenocytoid function in the establishment of anti-*Plasmodium* immunity.

In addition to PPO regulation, we also provide evidence that PGE2 signaling stimulates the production of AMPs in *An. gambiae*, similar to previous studies in other systems (4, 42). In light of this result, we demonstrate the role of PGE2 priming in mediating antibacterial immune responses that suppress bacterial growth. These effects are abrogated by *AgPGE2R*-silencing, indicating that *Ag*PGE2R is required for the antibacterial responses associated with PGE2 signaling.

In summary, our characterization of the prostaglandin receptor in *An. gambiae* provides important new insights into the roles of prostaglandin signaling in oenocytoid immune cell function and mosquito innate immunity. We demonstrate that *Ag*PGE2R mediates PPO and AMP gene expression that limit *Plasmodium* oocyst survivial and suppress bacterial infection, and establish that PGE2 promotes oenocytoid lysis. Together, this work reveals an integral role of oenocytoids in prostaglandin signaling and mosquito innate immunity.

## Materials and Methods

### Mosquito rearing and *Plasmodium* infection

*Anopheles gambiae* mosquitoes (Keele strain) were reared at 27°C and 80% relative humidity, with a 14/10 hour day/night cycle. Larvae were fed on fish flakes (Tetramin, Tetra), and adult mosquitoes were maintained on 10% sucrose solution. Female Swiss Webster mice were infected with a mCherry strain of *Plasmodium berghei* as previously described (16).

### Sequencing and phylogenetic analysis of the *Ag*PGE2 receptor

A putative prostaglandin E2 receptor (AGAP001561) was identified in *Anopheles gambiae* by BLAST P using orthologous sequence from the functionally characterized PGE2R in *M. sexta* (8). Cloning and sequencing of the full ORF cDNA of *Ag*PGE2R was performed from total RNA isolated from hemocytes perfused from non-blood fed female mosquitoes (n=50) using Direct-zol RNA Miniprep kit (Zymo research). After DNase I treatment following manufacture’s protocol (New England Biolabs), 200 ng of total RNA was used for cDNA synthesis using the RevertAid First Strand cDNA Synthesis kit (Thermo Fisher Scientific). To obtain full length ORF cDNA (1463 bp), PCR was performed at 96°C for 2 min, followed by 40 cycles of 96°C 30s, 62°C for 60s, 72°C 60s, with a final extension at 72°C for 5 min using pge2R-F and pge2R-R primers listed in Table S1. Following gel purification of the PCR product, the amplified PCR product was cloned into pJET1.2/blunt Cloning Vector (Thermo Fisher Scientific) and sequenced by the Iowa State DNA Core Facility. A phylogenetic tree was generated from amino acid sequences of human prostanoid and leukotriene receptors, and chemokine receptor 3 obtained from NCBI as previously described (8). Putative insect PGE2 receptors were predicted by protein BLAST search using PGE2 receptors sequenced from *M. sexta* and *An. gambiae*. The phylogenetic tree was produced with MEGA 7 software (43)

### *Ag*PGE2R expression analysis

Naïve and *P. berghei* infected mice were used for mosquito blood feeding and *Plasmodium* infection, respectively. To quantify relative abundance of receptor transcript level in various tissues, midgut, fat body, malpighian tubules and ovary were isolated from naïve (3-5 days old), 24 h blood-fed or 24 h *P. berghei* infected mosquitoes (n=40 per condition) in 1x sterile PBS. Hemolymph was separately perfused from mosquitoes (n=50) from similar conditions as described previously (16). Total RNA from isolated tissues was prepared using TRIzol (Thermo Fisher Scientific). Following hemolymph lysis in TRIzol reagent, total RNA was isolated using Direct-zol™RNA MiniPrep (Zymo Research). After DNase I treatment according to manufacture’s protocol (New England Biolabs), Total RNA from tissues (2 µg) and hemolymph (200 ng) was used for cDNA synthesis using the RevertAid First Strand cDNA Synthesis kit (Thermo Fisher Scientific). qRT-PCR was performed using PowerUp™SYBR®Green Master Mix (Thermo Fisher Scientific) with the ribosomal S7 protein transcript serving as an internal reference as previously (16). cDNA (1:5 dilution) amplification was performed with 500 nM of each specific primer pair using the following cycling conditions: 95°C for 10 min, 40 cycles with 95°C for 15 s and 65°C for 60 s. A comparative C_T_ (2^−ΔΔCt^) method was employed to evaluate relative transcript abundance for each transcript (44). A list of primers used for gene expression analyses are listed in Table S1.

### Western blot analysis

Western blot analysis was performed as previously described (16). Hemolymph was perfused from individual mosquitoes (n=35) at naïve, 24 h blood fed, or 24 h *P. berghei*-infected mosquitoes using incomplete buffer (anticoagulant solution without fetal bovine serum) containing a protease inhibitor cocktail (Sigma). Hemolymph protein concentrations were measured using Quick Start™Bradford Dye reagent (Bio-Rad). Protein samples (2 µg) were mixed with Bolt™LDS sampling buffer and sample reducing agent (Life Technologies), and heated at 70°C for 5 min before separation on 4-12% Bis-Tris Plus ready gel (Thermo Fisher Scientific). To determine PGE2R glycosylation in the hemolymph samples, protein samples were treated with PNGase F (Promega) according to manufacturer’s instruction under denaturing conditions. Samples were resolved using Bolt™MES SDS running buffer (Thermo Fisher Scientific) for 90 min at 100 V. Proteins were transferred to PVDF membrane in Bolt™Transfer buffer (Life Technologies) for 1 h at 20 V, and then blocked in TBST buffer (10 mM Tris base, 140 mM NaCl, 0.05% Tween 20, pH 7.6) containing 5% non-fat milk for 1 hour at RT. For western blotting, the membrane was incubated with a 1:1000 dilution of rabbit anti-*Ag*PGE2R (Cys-NRSMSQTPKSSSFTDSNIIR: third intracellular loop) affinity purified antibodies (3.1 mg/ml; Pacific Immunology), pre-immune serum (1:1000), or rabbit anti-serpin3 (SRPN3) antibodies (1:1000) (16) in TBST blocking buffer overnight at 4°C. Membranes were washed three times for 5 min in TBST, then incubated with a secondary anti-rabbit alkaline phosphatase-conjugated antibody (1:7500, Thermo Fisher Scientific) for 2 h at RT. Following washing in TBST, the membrane was incubated with 1-Step™NBT/BCIP (Thermo Fisher Scientific) to enable colorimetric detection.

### Immunofluorescence assays (IFAs)

Hemocyte immunofluorescence assays (IFAs) were performed as previously described (16). Hemolymph perfused from mosquitoes at naïve, 24 h post-blood meal and 24 h post-infection was placed on a multi-well glass slide (MP Biomedicals) and allowed to adhere at RT for 30 min. Cells were fixed with 4% paraformaldehyde for 15 minutes at RT, then washed three times in 1xPBS. Samples were incubated with blocking buffer (0.1% Triton X-100, 1% BSA in 1xPBS) for 24 h at 4°C and incubated with a 1:500 of rabbit anti-*Ag*PGE2R (CVRYRSATEPID: second extracellular loop) affinity purified antibodies (0.6 mg/ml) (Pacific Immunology), or pre-immune serum (1:1000) in blocking buffer overnight at 4°C. After washing 3 times in 1xPBS, an Alexa Fluor 568 goat anti-rabbit IgG (1:500, Thermo Fisher Scientific) secondary antibody was added in blocking buffer for 2 h at RT. Slides were rinsed three times in 1xPBS and mounted with ProLong®Diamond Antifade mountant with DAPI (Life Technologies). Images were analyzed by a fluorescence microscopy (Nikon Eclipse 50i, Nikon) and confocal microscopy (Leica SP5 X MP confocal/multiphoton microscope) at the Iowa State University Microscopy Facility.

### Phagocytic cell depletion using clodronate liposomes

To validate that the *Ag*PGE2R is predominantly expressed on non-phagocytic oenocytoid immune cells, female mosquitoes were treated with either control liposome or clodronate liposome as previously described to deplete phagocytic immune cell populations (16). Hemolymph was perfused from naive mosquitoes (n=40) at 24 h post-injection. Total RNA isolation, cDNA synthesis and qRT-PCR experiment were performed as described above. Relative abundance of *AgPGE2R* transcript level was determined between treatments.

### Endogenous PGE2 titer in the hemolymph

Mosquito samples for analysis of prostaglandin E2 (PGE2) were prepared as previously described (11). To determine how endogenous PGE2 level is regulated in the mosquito at different conditions (naïve, 24 h blood fed and 24 h *P. berghei* infection), mosquitoes (n=5) were homogenized in 400 µl of Hanks Buffer Salt Solution (HBSS, Sigma) with calcium and magnesium and incubated for 1 h at 28 °C. After centrifugation at 12,000 rpm for 10 minutes at 4°C, the supernatant was collected and stored at -80°C until analyzed. In addition to measurement of PGE2 titer in whole mosquito, hemolymph (5 µl per mosquito) was perfused from naïve, 24 h blood fed and 24 h *P. berghei* mosquitoes using the HBSS buffer, and a pool of hemolymph (50 µl) perfused from ten mosquitoes was used for measurement of PGE2 titer. PGE2 titer was measured by Prostaglandin E2 Monoclonal ELISA kit (Cayman Chemical) according to the manufacturer’s instructions. Absorbance was read at 412 nm using a microplate reader (Multi-mode reader, Biotek). PGE2 level was calculated using the Prostaglandin E2-Monoclonal program (4PL) at http://www.myassays.com (MyAssays LTd., UK).

### Silencing *Ag*PGE2R by RNAi

dsRNA synthesis was performed as previously described (16). The N-terminus of *AgPGE2R* including the 5’ UTR and first 61 amino acid residues was selected for dsRNA synthesis. Specific primers listed in Table S1 were designed to amplify a 421 bp DNA template for subsequent dsRNA production using cDNA synthesized from naïve whole female mosquitoes. The amplified PCR product was excised from an agarose gel, purified using Zymoclean Gel DNA Recovery Kit (Zymo research), and cloned into a pJET1.2/blunt vector (Thermo Fisher Scientific). The plasmid DNA was amplified with T7 primers (Table S1) at 96°C for 2 min, followed by initial 10 cycles of 96°C 30s, 58°C for 60s, 72°C 60s, and subsequent 30 cycles of 96°C 30s, 72°C for 60s, 72°C 60s, with a final extension at 72°C for 5 min. MEGAscript RNAi kit (Thermo Fisher scientific) was used for dsRNAs synthesis following the manufacturer’s instructions. dsRNA was precipitated with ethanol and resuspended in nuclease free water to 3 µg/µl. To determine the role of *Ag*PGE2R in *Plasmodium* development, mosquitoes (3-5 days old) were anesthetized on ice and intrathoracically injected with 69 nl (∼200 ng) of dsRNA per mosquito using Nanoject III injector (Drummond Scientific Company). The dsRNA treated mosquitoes were kept at 19°C for 4 days, then the effects of gene silencing on expression of immune effectors, development of *P. berghei*, and clearance of *E. coli* infection were evaluated. To determine RNAi efficiency, mosquitoes (n=15) at 4 days post-injection and mosquitoes (n=15) at 24 h post-*P. berghei* infection were collected for total RNA isolation, cDNA synthesis and qRT-PCR analysis as described above.

### Effects of PGE2 signaling on gene expression and PO activity

To determine if PGE2 signaling via *Ag*PGE2R influences prophenoloxidase (PPO) and antimicrobial peptide (AMP) gene expression as previously described (27), we examined phenoloxidase activity and gene expression following PGE2 priming. Naïve mosquitoes were injected with 69 nl of either 500 nM or 2 µM PGE2, with 0.05% ethanol in 1xPBS as a control. At 24 h post-injection, mosquitoes (n=15 per treatment) were collected for RNA isolation, cDNA synthesis, qRT-PCR analysis of *PPO* and *Ag*PGE2R gene expression and IFA as described above. At 24 h post-injection of either PGE2 or 0.05% ethanol PBS, a pool of hemolymph perfused from mosquitoes (n=15, 10 µl per mosquito) in nuclease free water was prepared for analysis of phenoloxidase (PO) activity. The perfused hemolymph (10 µl) was mixed with 90 µl of 3, 4-Dihydroxy-L-phenylalanine (L-DOPA, 4 mg/ml) dissolved in nuclease free water as previously described (16). After initial 10 min incubation at room temperature, PO activity was measured at 490 nm every 5 min for 30 min, then the final activity was measured at 60 min using a microplate reader. Similar experiments were performed following *AgPGE2R*-silencing 4 days post-injection of dsRNA. PO activity was measured in gene-silenced mosquitoes injected with 500 nM PGE2 using perfused hemolymph from mosquitoes (n=15) at 24 h post-injection.

### PGE2 priming on bacterial challenge

Bacterial challenge experiments were performed with slight modification from previous studies (45). Naïve female mosquitoes (3-5 day old) were intrathoracically injected with 69 nl of PGE2 (500 nM) or 0.05% ethanol PBS using a Nanoject III (Drummond Scientific). Kanamycin resistant *E*.*coli* constructed by transformation with mCherry2-N1 plasmid was cultured in Luria Bertani (LB) broth containing kanamycin (50 µg/ml) overnight at 37°C. This *E. coli* suspension (OD_600_ = 0.4, 10^8^ cells) was spun down at 8000 rpm for 5 min and the bacterial pellet was washed twice in 1xPBS. At 24 h post-PGE2 application, mosquitoes were challenged *E. coli* (∼6950 cells in 69 nl) and collected at 6 h and 24 h post infection. Individual mosquitoes were homogenized in 1 ml of LB broth. Mosquito homogenates (100 µl) were spread onto LB-kanamycin (50 µg/ml) agar plates. Bacterial plates were incubated overnight at 37°C and colony-forming units (CFU) per plate were assessed to quantify the level of infection. Additional experiments were performed in control- and *Ag*PGE2R-silenced mosquitoes to determine the role of *Ag*PGE2R in antimicrobial immunity was performed by similar *E. coli* challenge experiments as described above.

### Contributions of PGE2 and *Ag*PGE2R to *Plasmodium* survival

Naïve mosquitoes (3-5 days old) were injected with either 69 nl of 500 nM PGE2 or 0.05% ethanol PBS. At 24 h post-injection, mosquitoes were challenged with *P. berghei* infection and kept at 19°C. Oocyst survival was assessed at 8 days post-infection. To evaluate effect of silencing *Ag*PGE2R on parasite development, dsRNA treated mosquitoes were challenged with *P. berghei* and kept at 19°C until assessment of oocyst survival at either 2 days or 8 days post-infection. To determine whether anti-*Plasmodium* immunity mediated by PGE2 priming requires *Ag*PGE2R activation, mosquitoes with silenced PGE2R were primed with PGE2 (500 nM) before challenging with *P. berghei* infection and oocyst survival was assessed at 8 days post-infection.

### Effect of prophenoloxidases and antimicrobial peptides on *P. berghei* infection

Due to the effect of the PGE2 signaling system on regulation of a set of PPOs and antimicrobial peptide genes, RNAi experiments were carried out with selected genes: *PPO1* (AGAP002825), *PPO7* (AGAP004980), *PPO8* (AGAP004976) and *CEC1* (AGAP000693). T7 primers were designed using the E-RNAi web application (http://www.dkfz.de/signaling/e-rnai3/idseq.php) and listed in Table S1. dsRNA synthesis was performed as described above. The effects of gene silencing were measured 2 days post-injection in whole mosquitoes (n=15) by qRT-PCR as described above. Mosquitoes were challenged with *P. berghei* at 2 days post-injection of dsRNA, and oocyst survival was assessed at 7 days post-infection. The influence of PPOs in limiting oocyst survival was further examined in mosquitoes co-silenced with *PPO1* and *PPO3* at 2 days post-injection of dsRNA were primed with PGE2 (500 nM) and challenged with *P. berghei* infection. Oocyst survival was assessed at 7 days post-infection.

## Acknowledgments

This work was supported by a Postdoctoral Association Seed Grant Award from Iowa State University (to H.K.) and R21AI44705 from the National Institutes of Health, National Institute of Allergy and Infectious Diseases (to R.C.S.).

