## Supplementary material for "Identification of a prostaglandin E2 receptor that regulates mosquito oenocytoid immune cell function in limiting bacteria and parasite infection": Figures S1-S10, Table S1

### Supplemental Information

#### Supplemental Figures

**Figure S1.** Cloning of the *Ag*PGE2 receptor (*Ag*PGE2R) and amino acid translation.

**Figure S2.** Predicted secondary structure and phylogenetic analysis of the *An. gambiae* prostaglandin E2 receptor (AgPGE2R).

**Figure S3.** Relative abundance of *AgPGE2R* transcript in mosquito tissues under different physiological conditions.

**Figure S4**. *Ag*PGE2R localization to mosquito immune cells.

**Figure S5.** Analysis of mosquito PGE2 titers under different physiological conditions.

**Figure S6.** Efficiency of *AgPGE2R*-silencing.

**Figure S7.** PGE2 signaling promotes antimicrobial activity.

**Figure S8.** Evaluation of other PGE2-regulated genes by RNAi that do not result in oocyst phenotypes.

**Figure S9.** High concentrations of PGE2 that promote oenocytoid lysis result in increased PO activity.

**Figure S10.** Reduction in the expression of a subset of PPOs with 1 µM PGE2.

#### Supplemental Tables

**Table S1.** Primers for cloning, qRT-PCR, and dsRNA synthesis

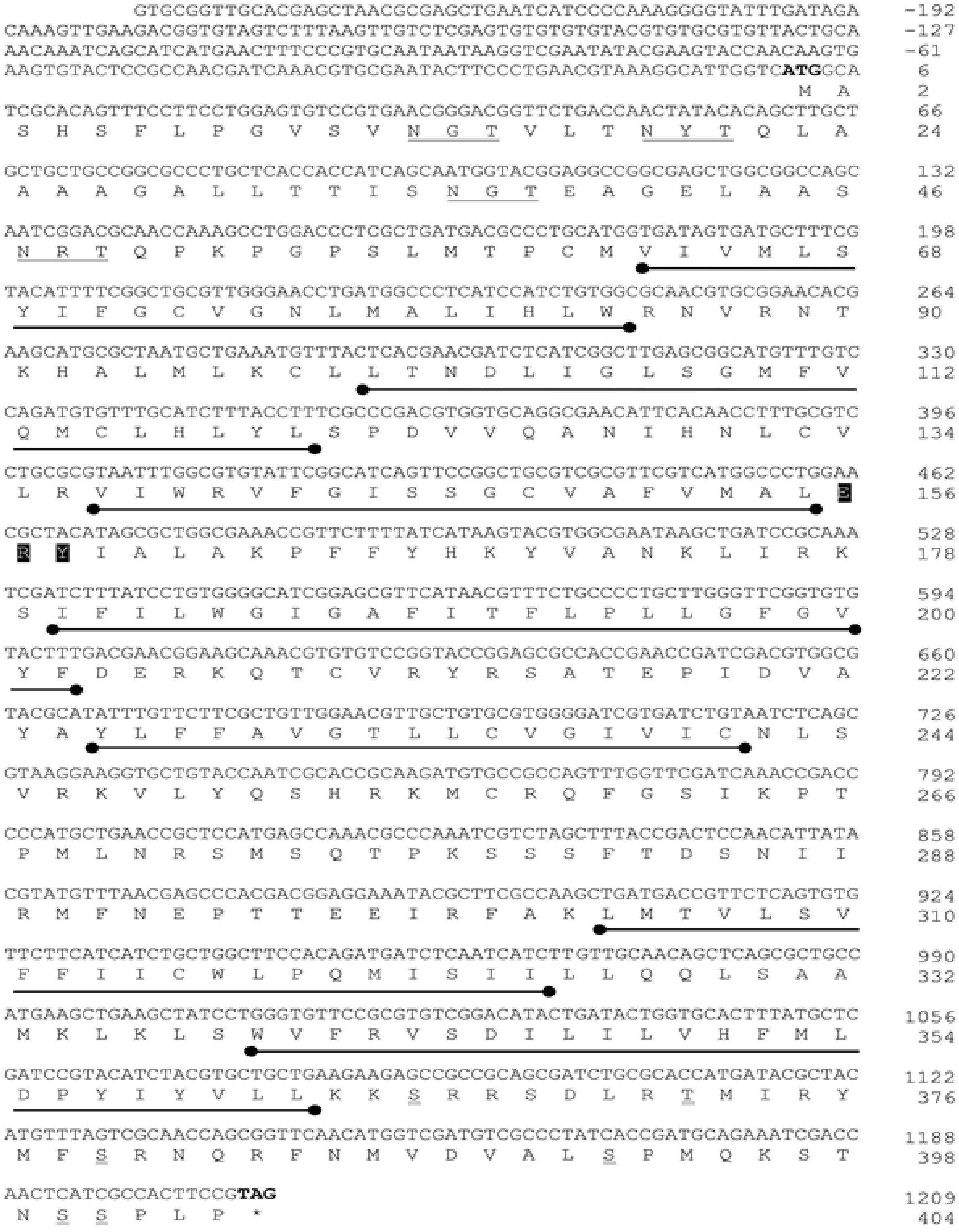

**Figure S1. Cloning of the *Ag*PGE2 receptor (*Ag*PGE2R) and amino acid translation.** cDNA prepared from perfused hemocytes of female adult naïve *An. gambiae* revealed a 1463 bp sequence, encoding a 404 amino acid residue protein. Seven transmembrane regions are underlined ( ●——● ), while highly conserved residues in Family A GPCRs (E156, R157, Y158) are shaded. Predicted N-linked glycosylation sites are underlined in the N-terminus. Single residues with predicted phosphorylation sites by protein-kinase A and C are underlined in the C-terminus.

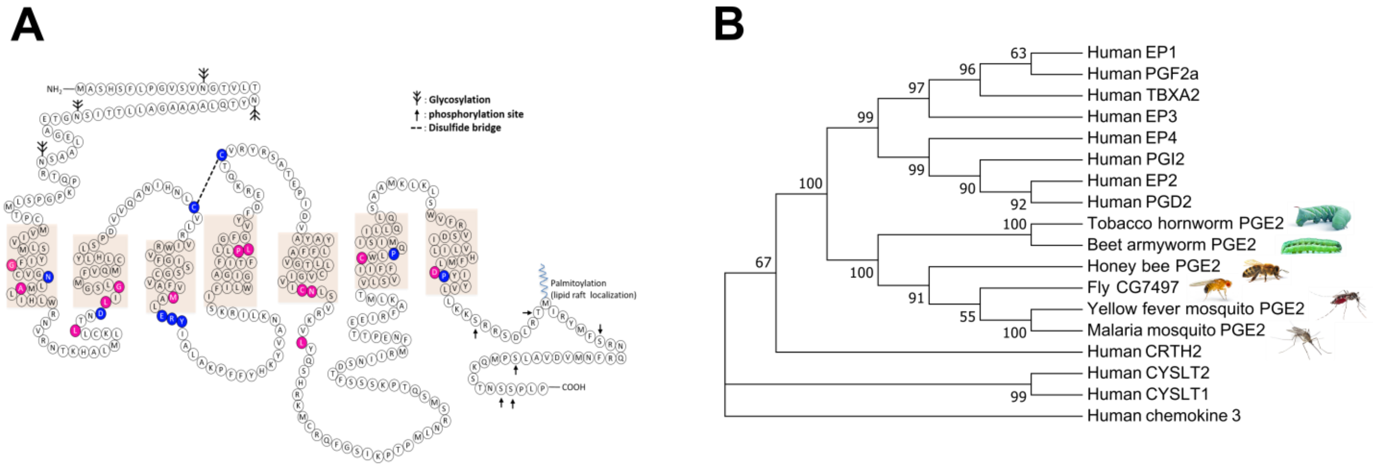

**Figure S2. Predicted secondary structure and phylogenetic analysis of the *An. gambiae* prostaglandin E2 receptor (*Ag*PGE2R)**. *Ag*PGE2R amino acid residues circled in blue are conserved in the rhodopsin-like family (Family A) of GPCRs, while residues circled in magenta are conserved with human prostanoid receptors (**A**). The seven transmembrane domains predicted by TMpred are displayed by beige-shaded boxes. Four N-glycosylated sites are predicted at the N-terminus and phosphorylation sites are located at the C-terminus. A single palmitoylation site was predicted by the presence of three cysteines at the C-terminus. (**B**) Phylogenetic analysis was performed with human prostanoid receptors, human leukotriene receptors, and putative insect PGE2 receptors (PGE2Rs). Human chemokine receptor 3 was used as outgroup to root the tree. Neighbor-joining method with 1000 bootstrap replicates was used to construct the tree using MEGA 7.0. The following GenBank accession number indicates different amino acid sequence found insects. Tobacco hornworm PGE2R (QDO79407.1), beet armyworm PGE2R (QEN91980.1), honey bee PGE2R (XP_001120030.2), fruit fly (NP_648999.2), yellow fever mosquito PGE2R (XP_021705923.1).

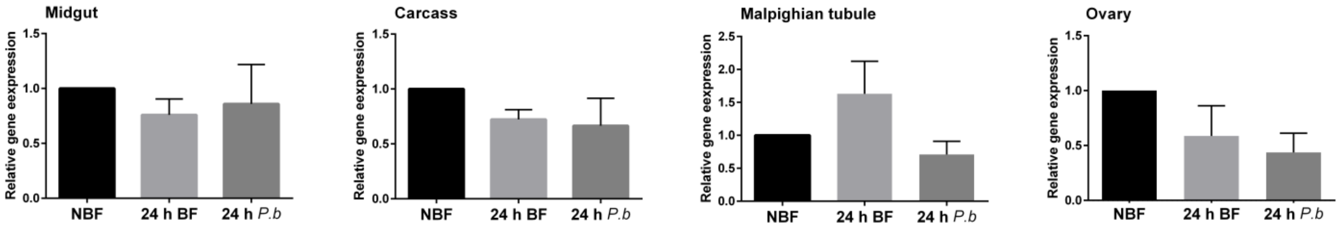

**Figure S3. Relative abundance of *AgPGE2R* transcript in mosquito tissues under different physiological conditions**. *AgPGE2R* expression was examined by qRT-PCR in midgut, fat body, ovary, and malphigian tubules under naïve (NBF), blood-fed (24 h BF), or *P. berghei*-infected (24 h *P.b*) conditions. When analyzed by a one-way ANOVA followed by a Tukey's multiple comparison test using GraphPad Prism 6.0, no differences in expression were detected across physiological conditions. Bars represent the mean ± SE of either three or four independent biological replicates.

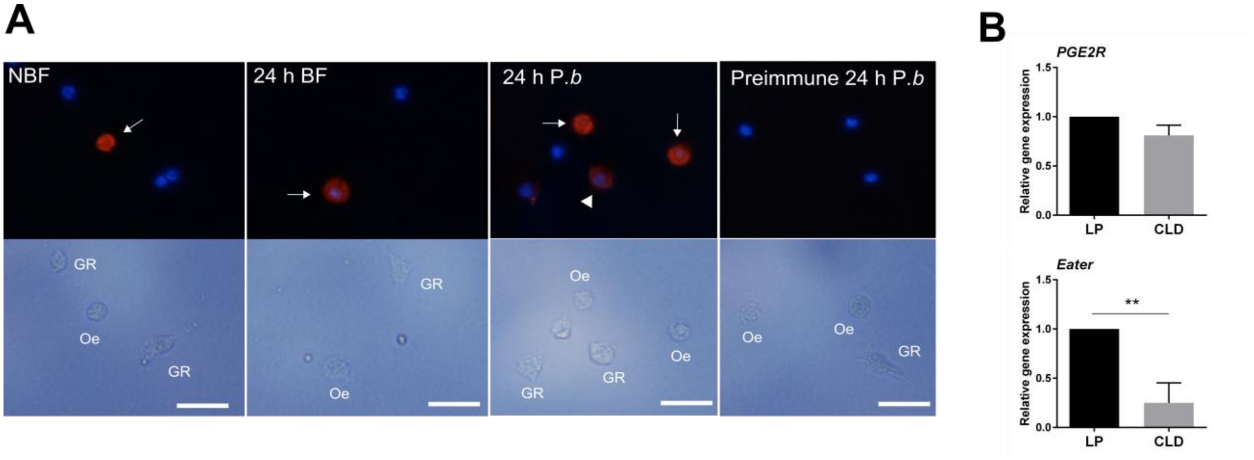

**Figure S4. *Ag*PGE2R localization to mosquito immune cells**. Immunofluorescence assays using an *Ag*PGE2R antibody recognizing extracellular loop 2 (ECL2) reveal that *Ag*PGE2R is predominantly expressed in oenocytoids (Oe, white arrow), although weak signal was detected in a subset of presumed granulocyte populations (Gr, white arrow head in 24 h *P.b*) (**A**). Scale bar represents 10 µm. (**B**) To validate that receptor signal is derived from the non-phagocytic oenocytoids, mosquitoes were treated with clodronate liposomes (CLD) to deplete phagocytic granulocytes. Empty liposomes (LP) were used as a control. qRT-PCR analysis demonstrates that *AgPGE2R* transcript is not reduced following CLD treatment, while the expression of the phagocyte marker *eater* was significantly reduced, supporting *AgPGE2R* is expressed primarily in the oenocytoids. Data were analyzed by an unpaired t using GraphPad Prism 6.0. Bars represent mean ± SE of three independent replications. Asterisks denote significance (***P* < 0.01).

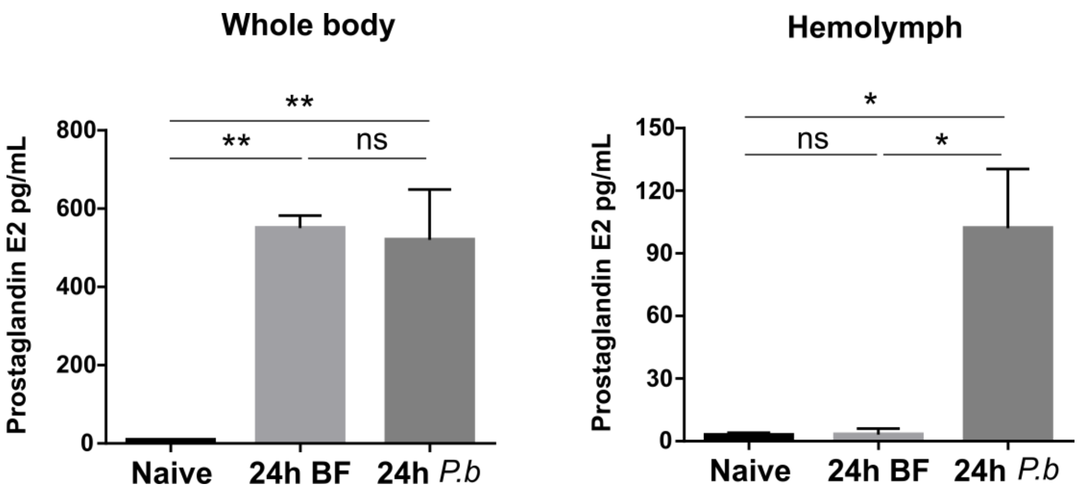

**Figure S5. Analysis of mosquito PGE2 titers under different physiological conditions**. PGE2 titers were measured from either whole mosquitoes or perfused hemolymph under naïve, at 24 h blood feeding (24 h BF), or 24h *P. berghei* infection (24 h *P.b*) conditions. For whole mosquitoes, samples were prepared from five mosquitoes, while hemolymph was collected from 10 mosquitoes per treatment. Data were analyzed using a one-way ANOVA followed by Tukey post-test using GraphPad Prism 6.0. Bars represents mean ± SE of three independent biological replicates. Asterisks denote significance (*$P < 0.05$, **$P < 0.01$); ns, not significant.

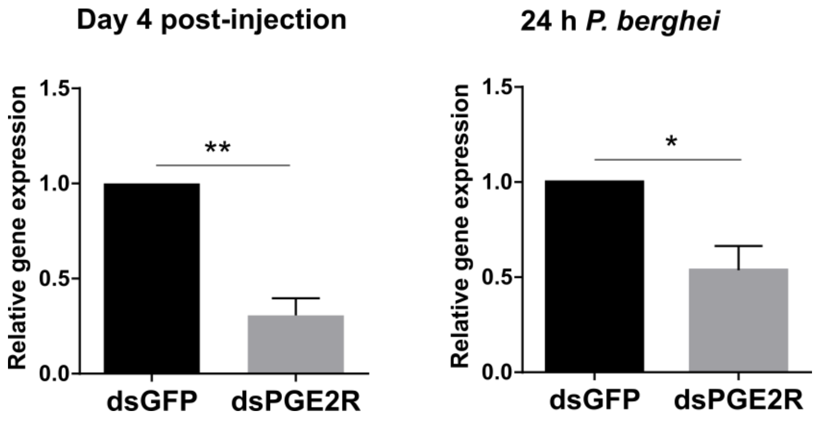

**Figure S6. Efficiency of *AgPGE2R*-silencing**. *AgPGE2R* expression was examined by qRT-PCR to validate the efficacy of RNAi. Whole mosquitoes were examined from naïve or 24h *P. berghei*-infected samples. Data were analyzed using an unpaired t-test to determine differences in *AgPGE2R* expression relative to GFP controls using GraphPad Prism 6.0. Bars represent mean ± SE of three independent biological replicates. Asterisks denote significance (**P* < 0.05, ***P* < 0.01).

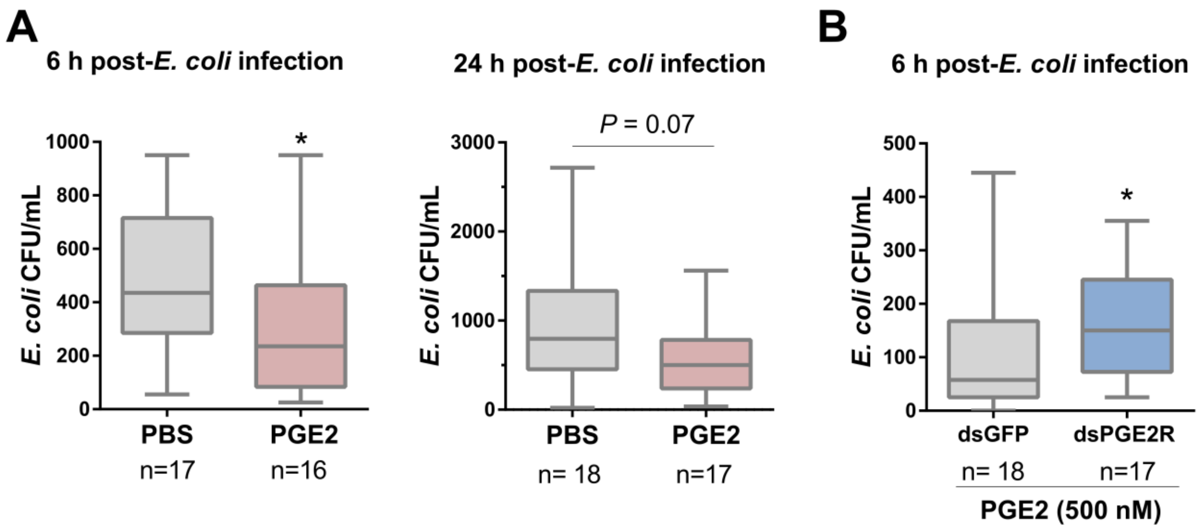

**Figure S7. PGE2 signaling promotes antimicrobial activity**. Naïve mosquitoes injected with 1x PBS or primed with PGE2 (500 nM) were challenged with *E. coli*, then bacteria titers were examined at 6 and 24 h post-infection (**A**). To validate that the antimicrobial activity of PGE2 occurred through *Ag*PGE2R, mosquitoes were silenced with dsGFP or dsPGE2R, primed with PGE2, then challenged with *E. coli* (**B**). *AgPGE2R*-silencing significantly impaired bacterial clearance compared to GFP control when evaluated 6 h post-infection. Data were analyzed by a Mann–Whitney test using GraphPad Prism 6.0. Bars represent mean ± SE of individual mosquitoes from three independent biological replicates. Asterisks denote significance (*$P < 0.05$).

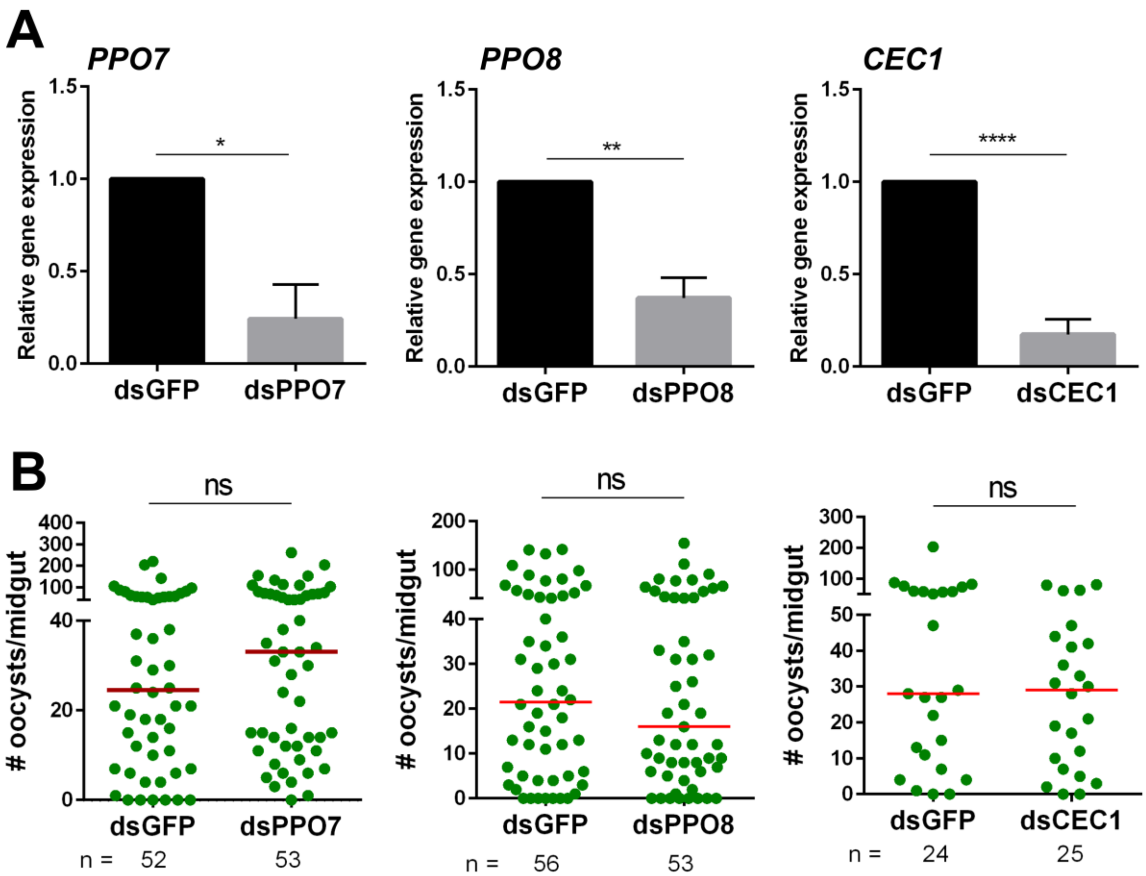

**Figure S8. Evaluation of other PGE2-regulated genes by RNAi that do not result in oocyst phenotypes.** The effects of dsRNA-mediated gene-silencing were evaluated in whole mosquitoes by qRT-PCR for *PPO7*, *PPO8*, and *CEC1* as compared to GFP controls (**A**). Data were analyzed by an unpaired t-test using GraphPad Prism 6.0. Bars represent mean ± SE of three independent biological replicates. (**B**) Additional experiments were performed for each knockdown to evaluate its contributions to malaria parasite survival. None of the examined genes influenced parasite survival when oocysts numbers were examined 7 days post-infection. Data were analyzed by a Mann–Whitney test to assess oocyst survival. Median oocyst numbers are indicated by the horizontal red line. Asterisks denote significance (**P* < 0.05, ***P* < 0.01, *****P* < 0.0001); ns, not significant.

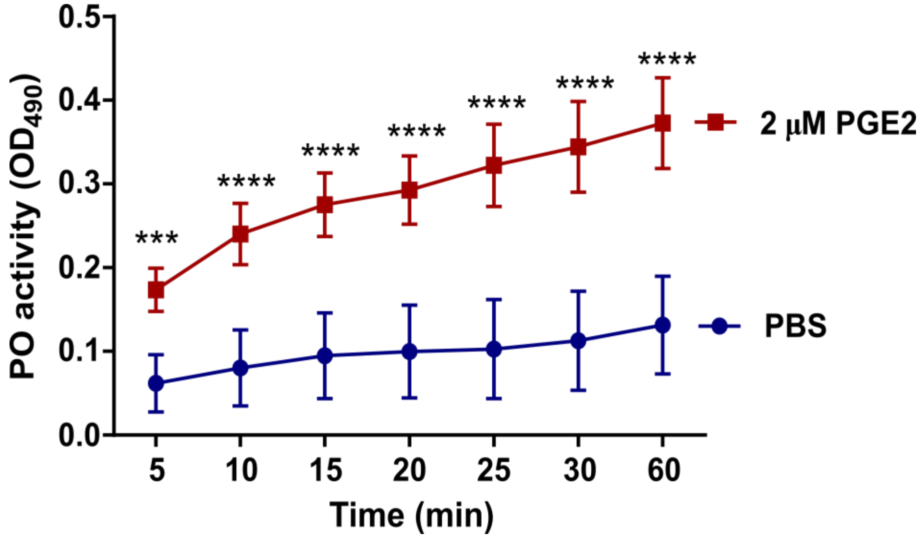

**Figure S9. High concentrations of PGE2 that promote oenocytoid lysis result in increased PO activity**. PO activity was measured from perfused hemolymph in mosquitoes primed with 2 μM PGE2 and compared to PBS controls 24 h post-treatment (n=15 per treatment). Measurements (OD490) were taken for DOPA conversion assays at 5-min intervals from 0 to 30 min, as well as a final readout at 60 min. Data were analyzed using a two-way repeated-measures ANOVA followed by Sidak's multiple comparisons using GraphPad Prism 6.0. Bars represent mean ± SE of 3 independent experiments. Asterisks denote significance (***$P$ < 0.001, ****$P$ < 0.0001).

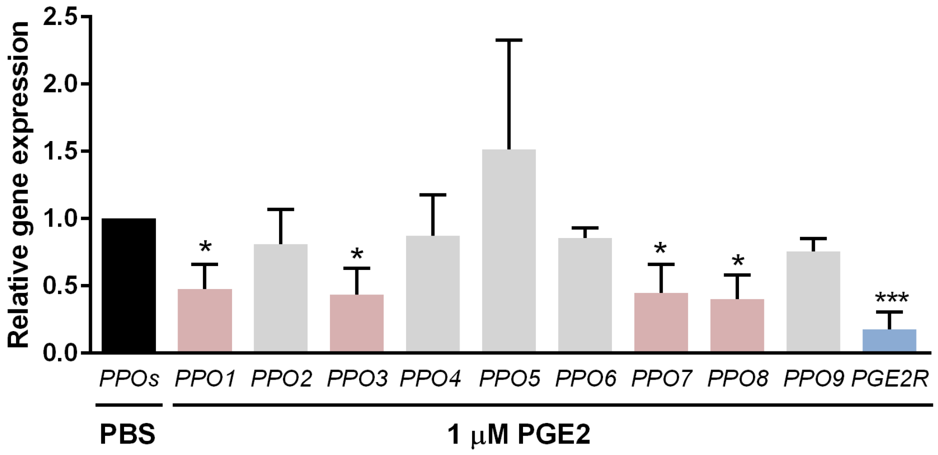

**Figure S10. Reduction in the expression of a subset of PPOs with 1 μM PGE2.** Similar to the effects 2 μM PGE2, the administration of 1 μM PGE2 promotes oenocytoid lysis reducing the expression of a subset of *PPOs* and *AgPGE2R*. Data were analyzed using an unpaired t test to determine differences in relative gene expression for each respective PPO gene between treatments using GraphPad Prism 6.0. Bars represent mean ± SE of three independent biological replicates. Asterisks denote significance (**P* < 0.05, ****P* < 0.001).

**Table S1. Primers for cloning, qRT-PCR, and dsRNA synthesis**

| Primer | Sequence (5'-3') | Gene ID |
|---|---|---|
| **For cloning** | | |
| pge2R-F | GTGCGGTTGCACGAGCTAAC | AGAP001561 |
| pge2R-R | CTACGGAAGTGGCGATGAGTTGG | |
| **For qRT-PCR** | | |
| pge2R-qF | GCTTCGCCAAGCTGATGACC | AGAP001561 |
| pge2R-qR | ACGCGGAACACCCAGGATAG | |
| ppo1-qF | GACTCTACCCGGATCGGAAG | AGAP002825 |
| ppo1-qR | ACTACCGTGATCGACTGGAC | |
| ppo2-qF | TTGCGATGGTGACCGATTTC | AGAP006258 |
| ppo2-qR | CGACGGTCCGGATACTTCTT | |
| ppo3-qF | CTATTCGCCATGATCTCCAACTACG | AGAP004975 |
| ppo3-qR | ATGACAGTGTTGGTGAAACGGATCT | |
| ppo4-qF | GCTACATACACGATCCGGACAACTC | AGAP004981 |
| ppo4-qR | CCACATCGTTAAATGCTAGCTCCTG | |
| ppo5-qF | GTTCTCCTGTCGCTATCCGA | AGAP012616 |
| ppo5-qR | CATTCGTCGCTTGAGCGTAT | |
| ppo6-qF | GCAGCGGTCACAGATTGATT | AGAP004977 |
| ppo6-qR | GCTCCGGTAGTGTTGTTCAC | |
| ppo7-qF | CAGCGATTGACGAAGGTGTT | AGAP004980 |
| ppo7-qR | GAAAGCAATACGTGCCCACT | |
| ppo8-qF | CCTTTGGTAACGTGGAGCAG | AGAP004976 |
| ppo8-qR | CTTCAAACCGCGAGACCATT | |
| ppo9-qF | TGTATCCATCTCGGACGCAA | AGAP004978 |
| ppo9-qR | AAGGTTGCCAACACGTTACC | |
| cec1-qF | CAGCAGAGAAGGCCCTACCG | AGAP000693 |
| cec1-qR | TCATGTTAGCAGAGCCGTCGT | |
| cec4-qF | CCACGCTGCTACTGTTCGGT | AGAP006722 |
| cec4-qR | CTGCAGTACGGGCACTACCT | |
| def1-qF | ATGCCGCGCTGGAGAACTAT | AGAP011294 |
| def1-qR | ATAGCGGCGAGCGATACAGT | |
| rps7-qF | ACCACCATCGAACACAAAGTTGACACT | AGAP010592 |
| rps7-qR | CTCCGATCTTTCACATTCCAGTAGCAC | |

**For dsRNA synthesis**

| | | |
|---|---|---|
| pge2R-T7F | TAATACGACTCACTATAGGGGTGCGGTTGCACGAGCTAAC | AGAP001561 |
| pge2R-T7R | TAATACGACTCACTATAGGGAGGGCGTCATCAGCGAGGGT | |
| ppo1-T7F | TAATACGACTCACTATAGGGCATCGGTTCCGAAATCCAG | AGAP002825 |
| ppo1-T7R | TAATACGACTCACTATAGGGCAGATCGAGATCGTGCGTAT | |
| ppo7-T7F | TAATACGACTCACTATAGGGGTACGAGTGCGGACACACC | AGAP004980 |
| ppo7-T7R | TAATACGACTCACTATAGGGGAATTTGAGGAAACGCTGCT | |
| ppo8-T7F | TAATACGACTCACTATAGGGAAAATTTGGCCAATCGCTTT | AGAP004976 |
| ppo8-T7R | TAATACGACTCACTATAGGGAACGAGTCCCGGCGTTTAAT | |
| cec1-T7F | TAATACGACTCACTATAGGGAGAGACCAACCAACCACCAA | AGAP000693 |
| cec1-T7R | TAATACGACTCACTATAGGGTTAGCAGAGCCGTCGTCTTA | |
| GFP-T7F | TTAATACGACTCACTATAGGGAGAATGGTGAGCAAGGGCGAGGAGCTGT | |
| GFP-T7R | TTAATACGACTCACTATAGGGAGATTACTTGTACAGCTCGTCCATGCC | |
